# Intracellular Delivery of Full-Length Antibodies via Poly-L-lysine-Coated PEG–PLGA Polymersomes Enables Non-invasive Pulmonary Immunotherapy

**DOI:** 10.1101/2025.09.16.676558

**Authors:** Nazar Vida, Naren Vyavahare, Jessica Larsen, Shivani Arora

## Abstract

The intracellular delivery of full-length antibodies holds transformative potential for treating a range of diseases but is hindered by poor cellular uptake, extracellular degradation, and limited encapsulation strategies. Here, we report a non-invasive, scalable, and biocompatible nanocarrier system using poly-L-lysine (PLL)-coated PEG–PLGA polymersomes to encapsulate and deliver full-length antibodies intracellularly via passive diffusion. Coating with 30 kDa ε-poly-L-lysine enhanced antibody loading efficiency by modulating membrane permeability and reduced vesicle surface charge, facilitating endocytic uptake while preserving antibody activity. The resulting polymersomes exhibited no detectable cytotoxicity and enabled efficient aerosol-based pulmonary delivery to lung epithelial cells in vitro. To our knowledge, this is the first demonstration of using PLL-coated PEG–PLGA polymersomes for passive intracellular delivery of full-length antibodies. This platform opens new avenues for lung-targeted immunotherapy and broadens the translational potential of intracellular antibody therapeutics.

**Graphical Abstract:** 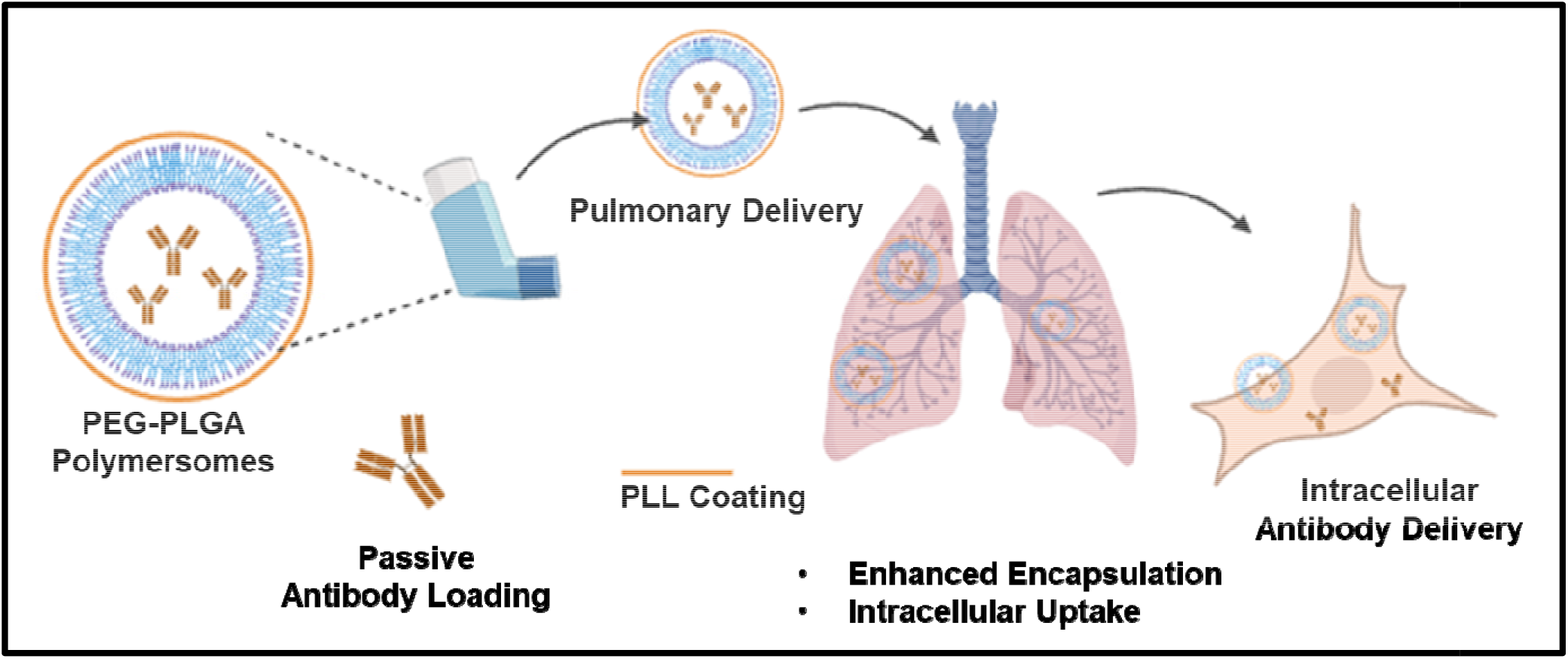

**Highlights:**

- PLL coating enhances passive antibody encapsulation efficiency
- PLL-coated polymersomes exhibit tunable, biocompatible surface charge profiles
- Efficient intracellular delivery of full-length antibodies via aerosol-mediated uptake in pulmonary epithelial cells
- Functional preservation of antibodies post-delivery validates PLL–PEG–PLGA as a protective nanocarrier

## Introduction

The intracellular delivery of antibodies holds immense potential across a variety of biomedical applications, including bone regeneration[1], vaccine development[2], cancer chemotherapy [3], and treatment of inflammatory disorders such as inflammatory bowel disease (IBD) [4]. However, the clinical translation of antibody-based intracellular therapies remains challenging due to several barriers. Native antibodies are susceptible to degradation, denaturation, aggregation, and hydrolysis in the extracellular environment. Moreover, their large size and hydrophilic nature hinder cellular uptake, and unprotected delivery may trigger immune responses or rapid clearance [5].

Encapsulation technologies offer a promising solution to these challenges by enabling controlled, stable, and targeted delivery of biologics such as antibodies [6]. For instance, Shrestha et al. demonstrated successful intracellular delivery of encapsulated Fab fragments of TNF-α antibodies for IBD therapy [4]. Despite this progress, the intracellular delivery of full-length antibodies (∼150 kDa) remains largely unexplored due to their size and structural complexity, which limit passive diffusion and membrane penetration.

Polymeric vesicles, or polymersomes (PSs), have emerged as adaptable nanocarriers for intracellular drug delivery. Their amphiphilic bilayer architecture—comprising a hydrophobic membrane and an aqueous core—enables the encapsulation of both hydrophobic and hydrophilic molecules, including therapeutic proteins and antibodies [7]. Among these, PEGylated polymosomes composed of polyethylene glycol (PEG) and poly(d,l-lactide-co-glycolide) (PLGA) stand out due to their established biocompatibility, tunable degradation profiles, and FDA-approved status of PLGA [8], [9], [10]. Moreover, PEG–PLGA PSs exhibit stealth properties, prolonged circulation time, and potential for surface engineering, making them attractive candidates for intracellular biologic delivery.

Despite these advantages, efficient encapsulation of full-length antibodies within PEG–PLGA polymersomes remains technically challenging. Passive loading of such large macromolecules is often limited by the low membrane permeability and dense polymeric structure of PSs. Consequently, active loading strategies or surface conjugation approaches are typically employed, albeit with increased complexity and potential for antibody denaturation.

To address these limitations, we explored a poly-L-lysine (PLL) coating strategy to enhance the loading and intracellular delivery potential of PEG–PLGA polymersomes. Rather than covalent surface functionalization, PLL was applied as an electrostatically driven coating, forming a cationic shell around the negatively charged PEG–PLGA vesicles. This non-covalent PLL coating enhances colloidal interactions, stabilizes vesicle morphology, and facilitates cellular uptake by promoting electrostatic interactions with negatively charged cell membranes [11], [12], [13]. Additionally, PLL coating can reduce the net negative charge of PEG–PLGA PSs, resulting in more favorable biological interactions [14], [15].

However, concerns about cationic polymer-induced toxicity persist. Ruenraroengsak et al. demonstrated that highly cationic nanoparticles may induce oxidative stress, mitochondrial dysfunction, and apoptosis in pulmonary cell types such as alveolar macrophages and epithelial cells [16]. These findings emphasize the importance of fine-tuning surface charge to maintain biocompatibility.

In this context, low to medium molecular weight ε-poly-L-lysine (e-PLL; 1–40 kDa) has gained attention as a safe, biodegradable, and food-grade cationic polymer. e-PLL is widely used as an antimicrobial preservative and has demonstrated excellent biocompatibility in biomedical settings, including pulmonary drug delivery applications[17], [18]. Its mucoadhesive and electrostatic properties make it particularly attractive for delivering macromolecules to the lungs.

Encouraged by these findings, we developed PEG–PLGA polymersomes coated with 40 kDa e-PLL to enable safe and efficient intracellular delivery of full-length antibodies. In this study, we demonstrate that PLL coating significantly enhances antibody encapsulation efficiency, likely by transiently increasing membrane permeability during the self-assembly process. Furthermore, the coated polymersomes exhibit robust cellular uptake, enable non-invasive pulmonary delivery via aerosolization, and preserve the functional activity of the encapsulated antibodies without inducing cytotoxicity.

To our knowledge, this is the first demonstration of using PLL-coated PEG–PLGA polymersomes to achieve passive loading and intracellular delivery of full-length antibodies, providing a scalable and biocompatible nanoplatform for lung-targeted immunotherapy and other biomedical applications.

## Results and Discussion

### Poly-L-lysine Coating Enhances Antibody Encapsulation in PEG–PLGA Polymersomes

To improve intracellular antibody delivery, we engineered polymersomes (PSs) composed of poly (ethylene glycol)-block-poly (lactic-co-glycolic acid) (PEG–PLGA) using a solvent injection method. PEG (2 kDa)-b-PLGA (4.5 kDa) was dissolved in DMSO (9.9% w/v) and injected at a controlled rate (5 μL/min) into an aqueous solution containing 2% w/v D-mannitol under stirring. The resulting PSs were lyophilized and stored under vacuum. Size distribution and morphology were characterized by dynamic light scattering (DLS) and transmission electron microscopy (TEM), respectively, confirming successful vesicle formation.

To enhance antibody encapsulation, we coated the PEG–PLGA PSs with poly-L-lysine (PLL) of varying molecular weights: 1 kDa and 30 kDa. Encapsulation of Alexa Fluor 488-conjugated anti-human IgG antibodies (SouthernBiotech, Cat# 2030-40) was assessed via flow cytometry by comparing the fluorescence signals of antibody-loaded versus blank PSs.

Blank (antibody-free) PSs, regardless of PLL coating, exhibited negligible fluorescence, confirming the absence of non-specific signal (**Figure 1**). In contrast, PSs encapsulating antibodies and coated with PLL (regardless of molecular weight) demonstrated a marked increase in fluorescence, indicating successful encapsulation. PLL-coated PSs prepared with 30 kDa PLL achieved an encapsulation efficiency of ∼80%, compared to only ∼10% for uncoated PSs. This dramatic enhancement suggests that PLL plays a dual role in promoting antibody retention and loading efficiency.

**Figure 1.**
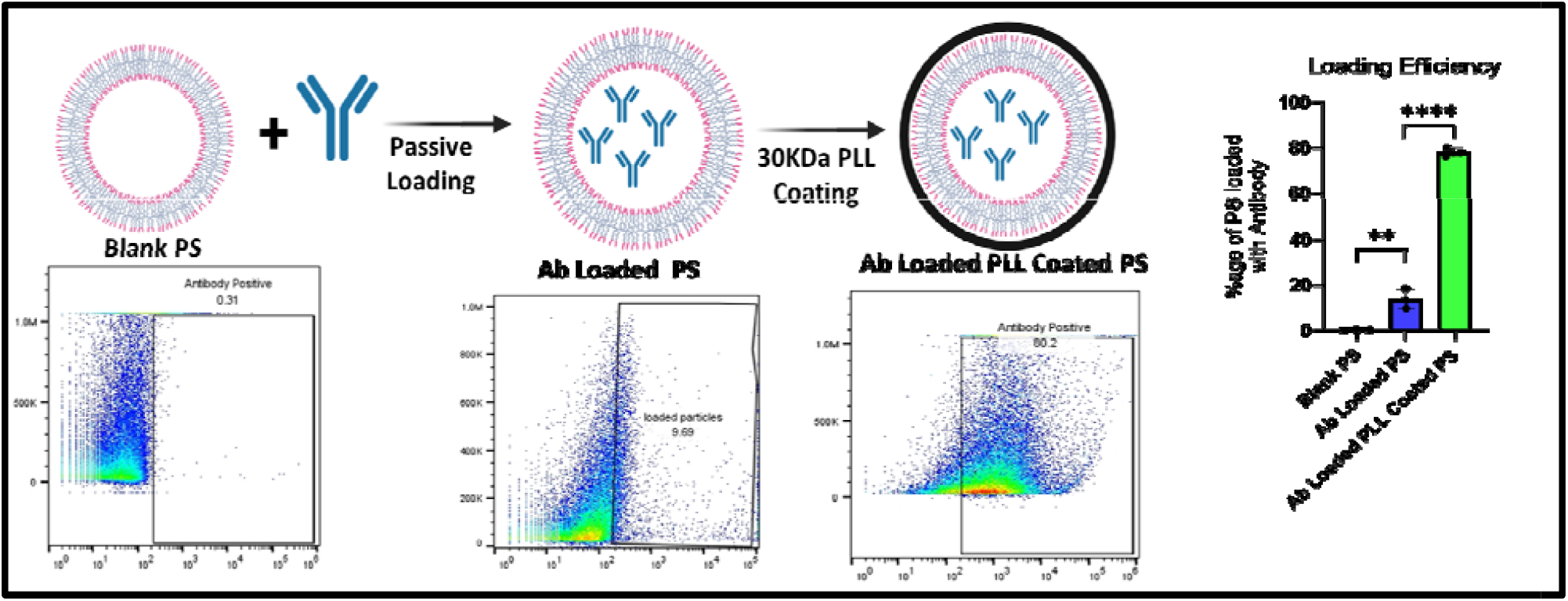
Synthesis and antibody loading of PLL-coated polymersomes (PLL-PS). **(A)** Schematic representation of the synthesis of antibody (Ab)-loaded PLL-PS. **(B)** Representative flow cytometry (FACS) plots of blank PS, Ab-loaded PS, and Ab-loaded PLL-PS. **(C)** Quantification of Ab loading efficiency by FACS. Results are represented as Mean ± SEM.

Two mechanisms may account for this observation: (1) PLL coating likely stabilizes the PS bilayer, reducing permeability and preventing diffusion-driven antibody loss; and (2) the positively charged PLL interacts electrostatically with the negatively charged antibody, enhancing local concentration during vesicle formation and favoring encapsulation. These interactions are particularly effective with higher molecular weight PLL, which may provide more extensive surface coverage and complexation capacity.

### PLL-coated polymersomes exhibit tunable, biocompatible surface charge profiles

Interestingly, 1 kDa PLL showed no difference in loading efficiency, while 30 kDa PLL optimized the balance between encapsulation efficiency and colloidal stability, as is validated by their **DLS and TEM** data (**Figure 2**). This finding aligns with previous studies indicating that moderately cationic coatings can improve cargo retention and delivery without inducing vesicle aggregation or cytotoxicity [1], [2].

**Figure 2.**
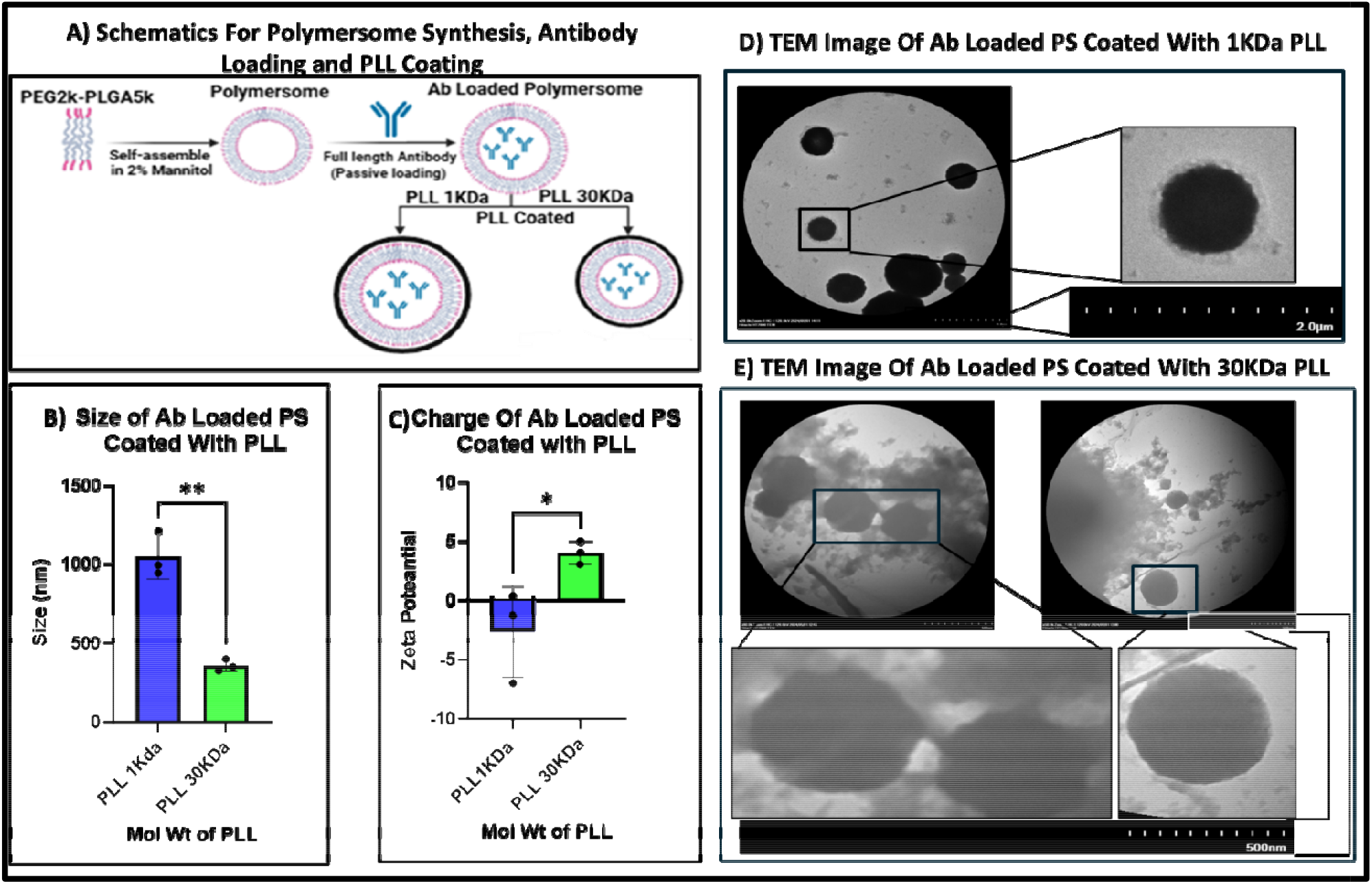
Characterization of antibody-loaded PLL-coated polymersomes (PLL-PS). **(A)** Schematic illustration of the synthesis of antibody-loaded PLL-PS. **(B–C)** Effect of PLL molecular weight on **(B)** polymersome size and **(C)** surface charge. **(D–E)** Representative TEM images of antibody-loaded polymersomes coated with **(D)** 1 kDa and **(E)** 30 kDa poly-L-lysine. Results represented as Mean ± SEM.

To assess the impact of poly-L-lysine (PLL) coating on surface charge characteristics of PEG– PLGA polymersomes (PSs), we analyzed zeta potential (ζ-potential) values of uncoated and PLL-coated PSs prepared using PLL of different molecular weights (1 kDa and 30 kDa). Uncoated PEG–PLGA PSs exhibited a strongly negative ζ-potential (∼−22 mV), consistent with the presence of free carboxylic acid termini on the PLGA moiety. This negative surface charge contributes to electrostatic repulsion from the anionic cell membrane, limiting cellular uptake efficiency.

Coating with low molecular weight PLL (1 kDa) partially neutralized this charge, yielding PSs with a moderately negative to near-neutral (-5 to-2Mv), whereas PS coated with 30KDa PLL displayed a ζ-potential of −5 mV to −8 mV. In contrast, PSs coated with 30 kDa PLL exhibited a positive ζ-potential (+ 4.5 mV, indicating successful electrostatic adsorption of the cationic polymer onto the PS surface and complete charge reversal. These results confirm that the surface charge of PEG-PLGA PSs can be systematically tuned by modulating PLL molecular weight and coating density.

Importantly, the ζ-potential values of the PLL-coated PEG–PLGA polymersomes remained within a physiologically acceptable range for pulmonary applications. Nanoparticles with surface charges between −10 mV and +15 mV have been reported to avoid significant cytotoxicity, hemolysis, or oxidative stress induction in mammalian systemsL[1], [2], [3]. This is particularly critical for pulmonary delivery, where excessive cationic surface charge is known to provoke adverse immune responses in lung-resident epithelial cells and alveolar macrophages [4].

Strongly cationic formulations are often used as mucoadhesive carriers, capitalizing on electrostatic interactions with the negatively charged mucosal lining of the trachea. However, this mucoadhesion can also restrict deeper penetration into the pulmonary tract. In contrast, near-neutral nanoparticles have been shown to bypass the mucosal barrier and facilitate deeper lung deposition, which is essential for efficient therapeutic delivery to alveolar regions[19].

### Efficient Intracellular Delivery of Antibody to Pulmonary Fibroblasts In Vitro

To evaluate the potential of PLL-coated PEG–PLGA polymersomes (PS–PLL) for intracellular antibody delivery, we performed in vitro uptake assays in primary human pulmonary fibroblasts.

First, we evaluated the loading capacity of the polymersomes to calculate the amount of antibody and PLL per mg of the PS. On average, we observed that 23ng of PLL is present per mg of PS and 27ng of Antibody is encapsulated by each mg of PS (Figure 3A)

**Figure 3.**
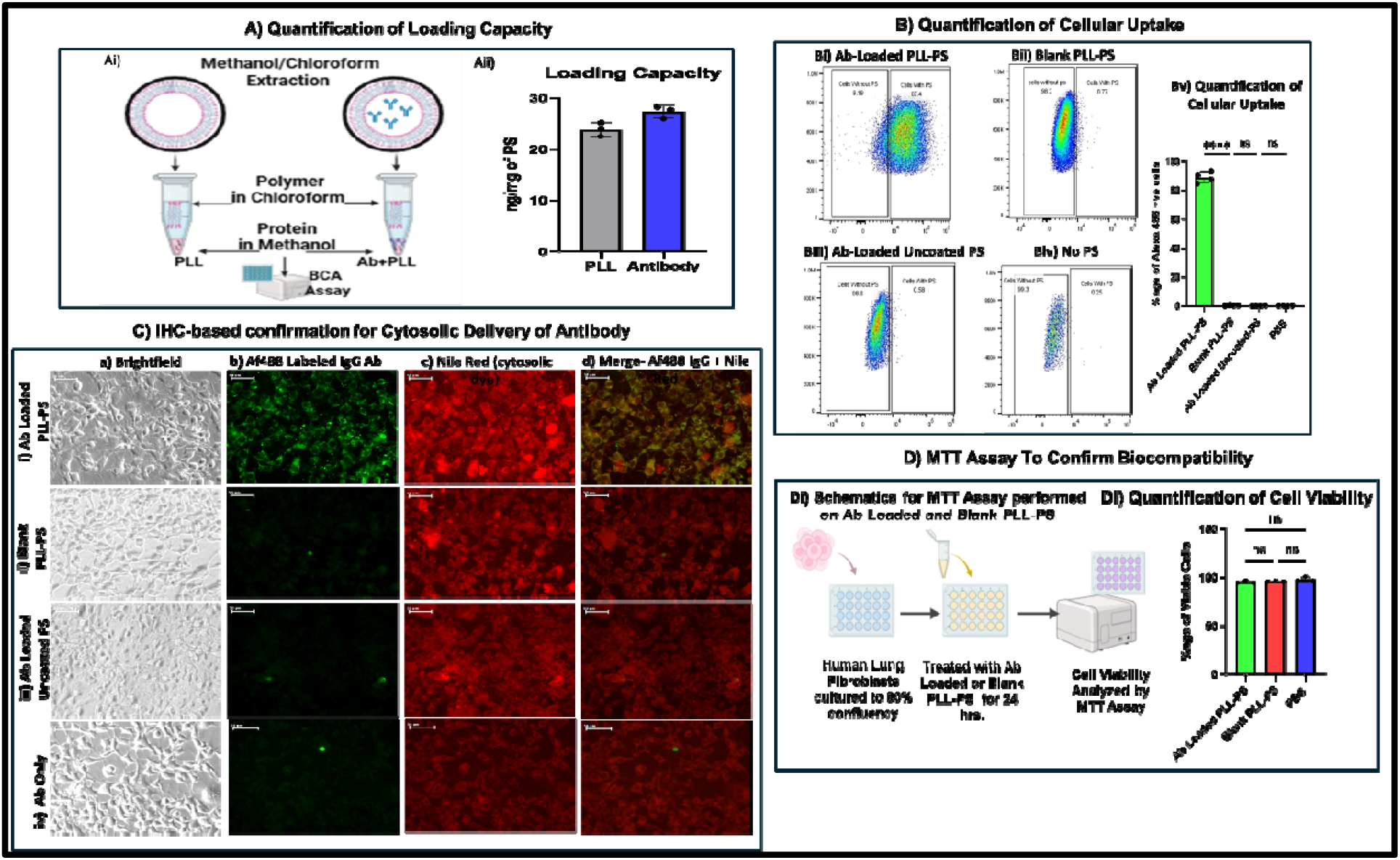
Evaluation of 30 kDa PLL-coated polymersomes (PLL-PS). **(A)** Antibody loading capacity and PLL content per mg of polymersomes. **(B–C)** Cellular uptake in primary human lung fibroblasts assessed by **(B)** flow cytometry and **(C)** immunohistochemistry. **(D)** Cytotoxicity analysis by MTT assay. Data represented as Mean ± SEM.

In Vitro Evaluation Using Human Pulmonary Fibroblasts-Primary human pulmonary fibroblasts were treated with AF488-labeled anti-human IgG either in free form or encapsulated in PLL (30 kDa)-coated polymersomes. Cells were exposed to the formulations overnight to (250ug of PS in culture media) and assessed for antibody uptake via flow cytometry and immunofluorescence microscopy. Untreated cells, blank PS–PLL, uncoated antibody-loaded PS, and free antibody served as controls.

Flow cytometry demonstrated a marked increase in fluorescence signals in cells treated with PLL-coated antibody-loaded PS, compared to cells treated with uncoated PS or free antibody (Figure 3B). This indicates enhanced cellular uptake and reduced extracellular retention when PS was coated with 30KDa PLL. Confocal microscopy confirmed cytosolic localization of the internalized antibody in PLL-coated PS–treated cells, as visualized by co-staining with Nile Red (Figure 3C).

Importantly, no signs of cytotoxicity, membrane damage, or altered cell morphology were observed under phase-contrast imaging (Figure 3a), and cell viability remained above 96% by MTT assay (Figure 3D). These findings confirm that PLL-coated PS provides a safe and efficient platform for intracellular antibody delivery in primary lung fibroblasts.

**To evaluate whether the antibody retains its functional integrity following encapsulation and intracellular delivery via PLL-coated PEG–PLGA polymersomes (PS–PLL)**, we employed a previously validated full-length monoclonal antibody targeting human NLRP3. This antibody, as demonstrated by Cliff et al., acts as a PROTAC-like degrader, reducing NLRP3 protein levels and downstream inflammasome activation.

Human pulmonary fibroblasts were cultured in fibroblast growth medium. Control conditions included cells treated with blank PS–PLL, free antibody (5µg), or uncoated antibody-loaded PS. After treatment, cells were washed and stimulated with 5 mM LPS for 6 hrs. and 10 µM nigericin for 45 minutes to induce NLRP3 inflammasome activation and IL-1β secretion. The cells were then treated with NLRP3 antibody-loaded PS–PLL (250 µg/well), NLRP3 antibody (5uM); Blank PLL-PS and media only.

Cell supernatants were collected, clarified by centrifugation, and analyzed using a human IL-1β– specific sandwich ELISA (Figure 4A). As anticipated, LPS/nigericin stimulation of untreated and control-treated cells resulted in robust IL-1β secretion. In contrast, fibroblasts pre-treated with antibody-loaded PS–PLL exhibited a marked reduction in IL-1β levels (Figure 4B), indicating successful intracellular delivery and preserved functional activity of the encapsulated antibody.

**Figure 4.**
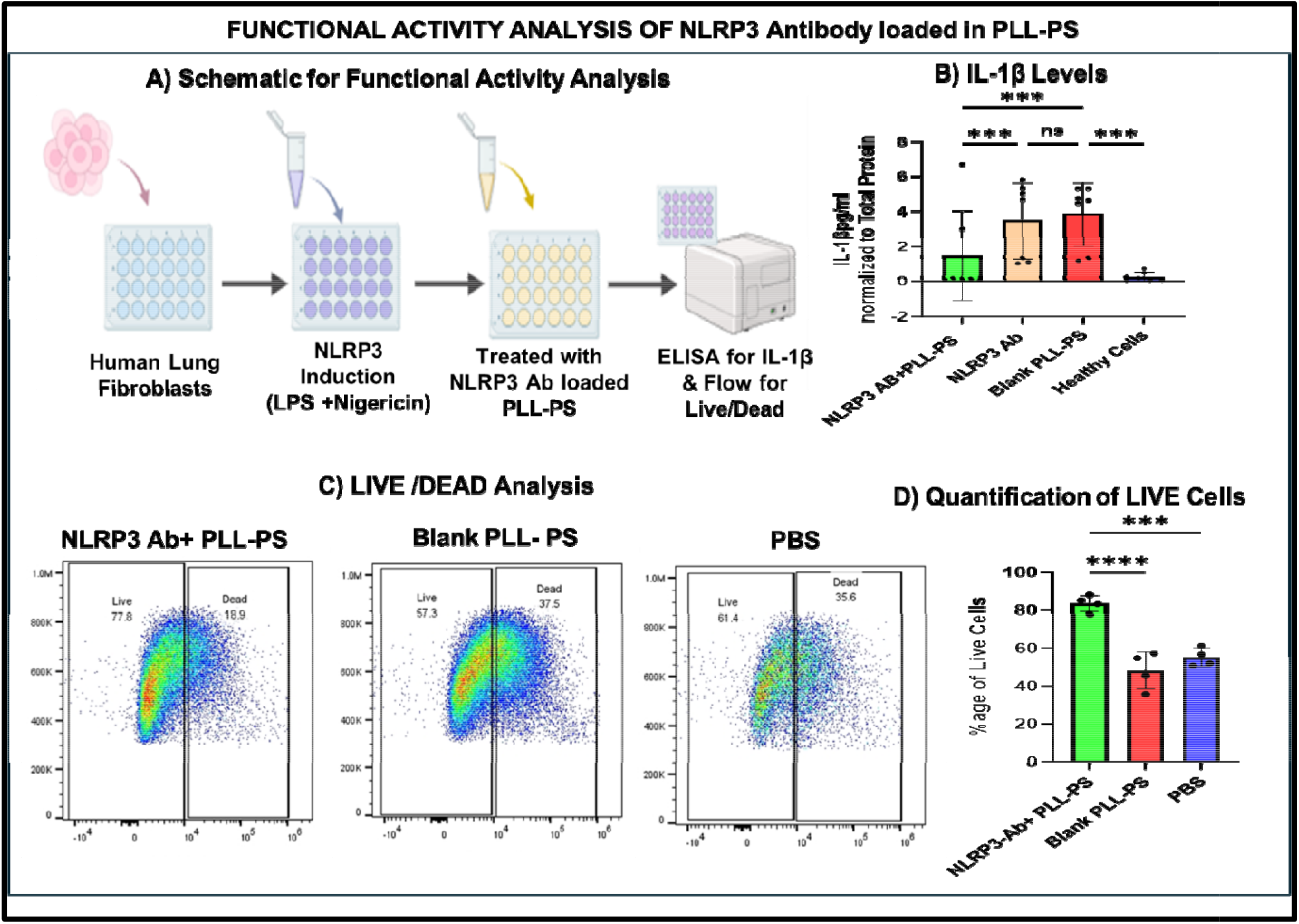
Functional activity analysis of NLRP3 antibody delivered using PLL-PS. **(A)** Schematic of experimental design using primary human lung fibroblasts. **(B)** Quantification of IL-1β levels in culture supernatants by ELISA, normalized to total protein. **(C)** Representative flow cytometry plots from Live/Dead assay following NLRP3 induction and treatment with NLRP3 Ab–loaded PLL-PS, blank PLL-PS, or untreated healthy cells. **(D)** Quantification of live cells across treatment groups. Data shown as Mean ± SEM.

Notably, neither free antibody nor uncoated antibody-loaded PS significantly attenuated IL-1β release, highlighting the critical role of PLL coating in enabling both mucosal penetration and efficient intracellular delivery. These results demonstrate that PS–PLL nanocarriers not only protect the antibody from degradation during formulation and aerosol delivery but also maintain its biological function within target cells.

Additionally, DAPI-based live/dead cell analysis revealed a significant increase in the percentage of live cells in the group treated with NLRP3 antibody–loaded PS–PLL compared to both untreated cells and those treated with blank PS–PLL (Figure 4C-D). This observation suggests that antibody-mediated inhibition of NLRP3 may confer cytoprotective effects, likely by reducing caspase-1–dependent pyroptotic or apoptotic cell death. While these findings are consistent with NLRP3 inhibition and downstream suppression of caspase-1 activation, further mechanistic validation through direct quantification of NLRP3 and caspase-1 protein levels is warranted.

### Efficient delivery of Antibody to Lung-Resident Cells In Vivo via Aerosolized PLL-Coated Polymersomes

In Vivo Aerosol-Mediated Pulmonary Delivery-To test the feasibility of aerosol-based antibody delivery to the lung, antibody-loaded PS–PLL were nebulized and delivered via nose-only inhalation to mice. After 2 hours, lung tissues were harvested and analyzed for intracellular antibody distribution. Confocal imaging of lung cryosections revealed a strong intracellular AF488 signal in alveolar and bronchial epithelial cells, indicating successful cellular uptake of the antibody delivered via aerosolized PS–PLL.

Control groups receiving saline showed minimal to no signal in lungs as measured by IVIS imaging, supporting the conclusion that PLL-coated PS can penetrate the mucosal barrier and facilitate cellular internalization (Figure 5A).

**Figure 5.**
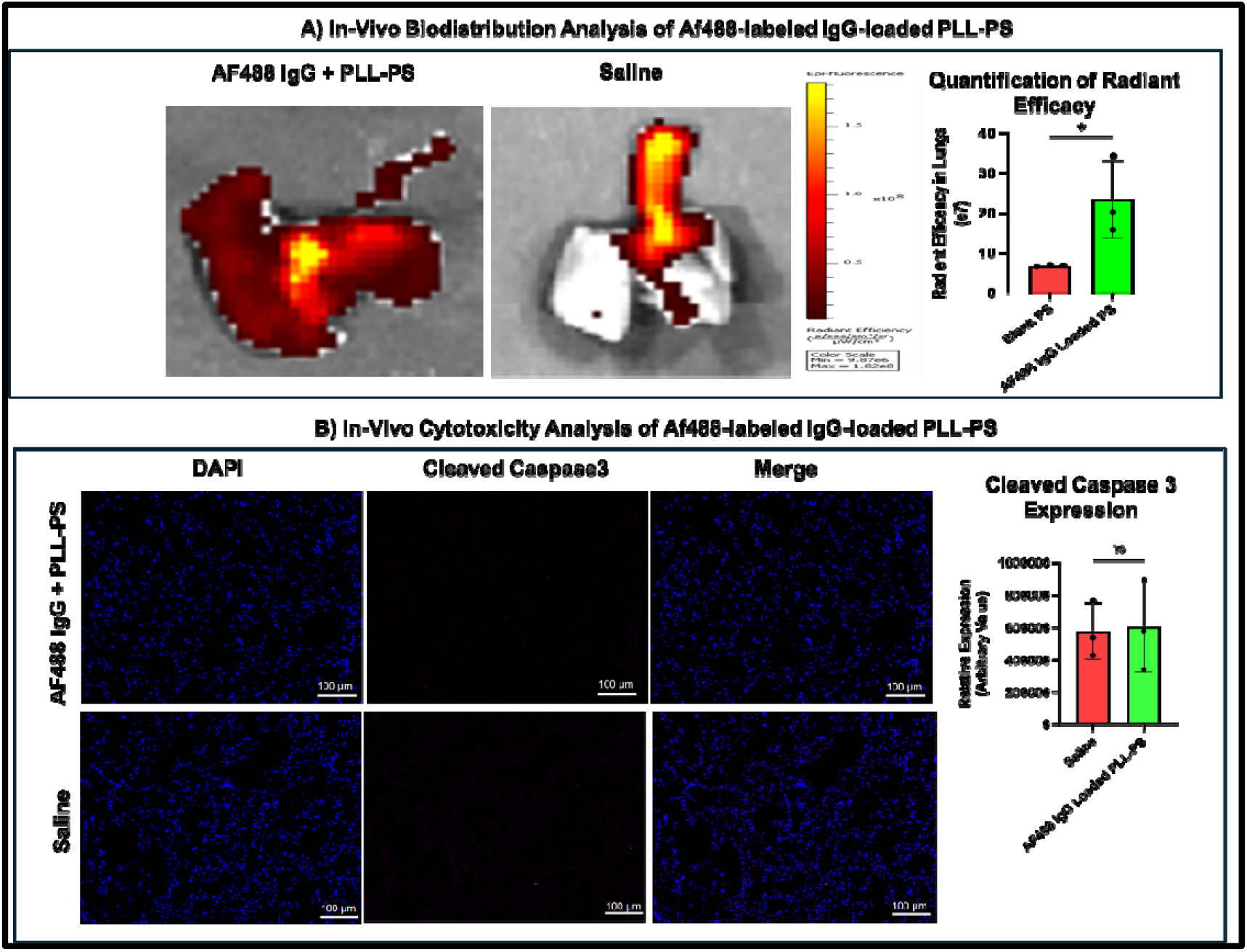
In vivo biodistribution following pulmonary delivery via aerosol pump. **(A)** IVIS images and IHC-stained lung sections from animals treated with AF488-labeled IgG antibody– loaded PLL-PS or saline. **(B)** In vivo cytotoxicity analysis of lungs treated with AF488-labeled IgG antibody–loaded PLL-PS or saline, assessed by IHC staining for cleaved Caspase-3 expression. Results represented as Mean ± SEM.

Furthermore, no evidence of acute cytotoxicity was observed, as indicated by the absence of significant Caspase-3 expression in lung sections, thereby demonstrating in vivo safety (Figure 5B). These findings affirm the biocompatibility of the 30 kDa PLL-coated PS formulation for pulmonary applications.

## Discussion

We demonstrated that the encapsulation efficiency of the antibody in polymersomes increased dramatically from 10% to 80% after coating the nanoparticle surface with PLL, consistent with other studies showing that coated polymersomes remain non-leaky and retain their payload for extended periods after coating a second layer to the structure[20], and as demonstrated by Coustet et al. [21]The layer-by-layer assembly approach produces a thicker shell and consequently increases loading capacity. A plausible mechanism underlying this improvement is that PLL coating stabilizes the polymersome bilayer, reducing membrane permeability and thereby preventing diffusion-driven antibody loss.

In addition, protein–antibody interactions have been shown to vary depending on environmental conditions. Robert et al. [22] reported that such interactions can take the form of short-range adhesive forces or longer-range electrostatic interactions. Building on this, we propose that the positively charged PLL interacts electrostatically with the negatively charged antibody. This interaction not only enhances the local antibody concentration during vesicle formation but also favors efficient encapsulation. These effects are particularly pronounced when higher molecular weight PLL is employed, as it can provide more extensive surface coverage and a greater complexation capacity with the antibody.

Based on our findings, the molecular weight of PLL significantly influences both particles’ size and zeta potential, as we observed that going from 1kDa to 30kDa PLL made the particle diameter smaller from approximately 1µm to 350nm and raised the zeta potential from-2 to +5 mV. As suggested by Volodkin et al. [23] Lower molecular weight PLL adsorbs flatly on negatively charged surfaces, while higher PLL 40kDa forms more loops and tails, causing greater overcharging, better stabilization and lower PDI of vesicles by reducing short-range attraction, hence smaller diameter[24]. High-molecular-weight PLL is therefore widely utilized in a range of practical biomedical applications [25].

The synthesized Polymersomes coated with PLL 30kDa showed no cytotoxicity in vitro. When administered in vivo, it also proved to be nontoxic, as no Caspase-3 expression was seen in treated mice. It is known that for pulmonary drug delivery, surface charge is especially critical. To avoid entrapment in the negatively charged mucus layer and to achieve deeper lung penetration, particles must exhibit near-neutral zeta potential during initial transport[26]. Near-neutral particles pass through the mucus layer faster due to a reduction in electrostatic and hydrophobic interactions[27].

Once beyond the mucus barrier, however, a shift toward a slightly positive potential is desirable, as this facilitates optimal interactions with the negatively charged cell membranes of lung epithelial and alveolar cells. A modestly positive zeta potential of around +5 mV is considered ideal [19]. At this charge range, the delivery vehicles are efficiently taken up by target cells while minimizing toxicity and avoiding excessive adhesion to the mucus layer. If the particles are too positively charged, they may bind strongly to mucosal components, become trapped, and be cleared rapidly via mucociliary mechanisms, thereby reducing therapeutic efficacy [28], [29].

## Conclusion

These studies collectively establish PLL-coated PEG–PLGA polymersomes as a robust, scalable, and biocompatible platform for the intracellular delivery of full-length therapeutic antibodies. By leveraging mid–molecular weight PLL (30KDa) for surface coating, we achieved precise modulation of nanoparticle surface charge (∼+5 mV), enabling efficient cellular internalization while maintaining safety in both in vitro and in vivo settings.

Importantly, this approach avoids the need for chemical modification of the antibody cargo, preserving its structural integrity and biological activity post-delivery. The functional suppression of NLRP3-mediated IL-1β secretion in pulmonary fibroblasts confirms that antibodies remain active following encapsulation, aerosolization, and intracellular uptake—validating PLL–PEG– PLGA polymersomes as protective nanocarriers.

The tunability of surface electrostatics via PLL coating provides a strategic advantage, allowing formulation customization based on the charge sensitivity of different target tissues or disease contexts. For pulmonary applications, the formulation’s ability to traverse the mucosal barrier and deliver biologics directly to lung-resident cells without inducing cytotoxicity positions it as a promising platform for inhaled antibody therapies.

Overall, our work highlights a non-covalent, modular delivery strategy that can be readily adapted to a broad range of intracellular protein therapeutics, offering a compelling solution to a long-standing barrier in biologic drug delivery.

## Materials and Methods

Poly (ethylene glycol) methyl ether-block-poly(lactide-co-glycolide), PEGL(2KDa)-b-PLGA (4.5 KDa) (Sigma-Aldrich cat# 764825). D-mannitol (Fisher Scientific, M120). Poly-L-lysine hydrobromide (PLL) 1Kda (Sigma-Aldrich cat# P0879). PLL 30KDa (Sigma-Aldrich, P2636); Anti-human IgG AF488 (SoutherenBiotech; 2030-40); Primary human Lung Fibroblasts (ATCC, PCS-201-013); Fibroblast Basal Media (ATCC, PCS-201-030); Fibroblast growth Kit (ATCC, PCS-201-041), NLRP3 Ab (Novus Biologicals; NBP2-12446); Anti mouse Caspase 3 antibody, Anti-Rabbit IgG Cy5 conjugated, MTT assay kit (Invitrogen, V13154), IL-1β Elisa Kit (Invitrogen, DY201-05), DAPI (Invitrogen, D 3571), BCA assay Kit (Thermo Fischer, 23227), Uranyl Acetate, TEM Grid, 300 kDa Float-A-Lyzer (Biotech CE Trial kit dialysis membrane).

### PS synthesis and characterization

PEG-PLGA PSs were formed using the solvent injection method[15], [30]. Briefly, PEG-PLGA was dissolved in DMSO at a concentration of 9.9 w/v%. The polymer solution was injected into 2 w/v% mannitol in water at a rate of 5 μL/min under stirring, using a 20-gauge needle and a syringe pump. PSs were lyophilized after slow freezing at-80°C and stored under vacuum conditions. The size distribution of PEG-PLGA PSs was obtained using Malvern Zetasizer Nano ZS (Malvern Ltd.) at 25°C. The morphology of PSs was observed using transmission electron microscope (TEM) imaging.

### TEM Imaging

The grid with PSs was stained using uranyl acetate and imaged with Hitachi HT7830 UHR 120LkV TEM (Tokyo, Japan).

### Loading Efficiency

The loading efficiency, defined as the percentage of PSs successfully encapsulating the antibody. Loading efficiency was determined using flow cytometry. PSs without antibody served as the negative control, while PS–antibody and PS–PLL–antibody samples were analyzed to quantify the proportion of particles loaded with antibody. L

### Loading capacity

To evaluate the loading capacity, antibody-loaded as well as blank PS were coated with PLL as described above. Briefly described-500µL of antibody (200 µg/mL) solution (for Ab-Loaded PS) or water (For blank PS) was added to 20 mg of PS. The mixture was then incubated with PLL 30KDa (200 μg) for 3 hrs. The antibody-loaded and blank PS–PLL particles were dialyzed against water using a 300 kDa Float-A-Lyzer for 24 hours at 4 °C, with the external solution replaced at 1, 2, 3, and 4 hours. After dialysis, the particles were freeze-dried. The resulting powder was weighed and resuspended in 100 µL of PBS. Next, we use the methanol/chloroform method to break PS and to extract the protein for quantification using the BCA method. Briefly, 400 µL of methanol was added to the PS solution obtained above and vortexed. This was followed by the addition of 100 µL of chloroform and vortexing. Subsequently, 300 µL of DI water was added, producing a cloudy suspension with precipitate. The mixture was centrifuged at 14,000 g for 1 minute. The top layer was carefully removed, and the methanol phase was discarded. The resulting pellet was air-dried for 5 minutes. Finally, the pellet was resuspended in 500 µL of 1M Tris-HCL buffer containing 10%SDS, and protein quantification was performed using the BCA assay [31]. Using this method, we were able to calculate the amount of PLL coating/mg of PS and the antibody Loading Capacity of PS (amount of antibody/mg of PS).

### Cellular uptake and Cytotoxicity Analysis of Antibody-loaded PLL-coated Polymosomes

**To** test the intracellular antibody delivery ability and to assess the cytotoxicity of the PLL (30KDa) coated PS, we performed in vitro cellular uptake analysis of the final formulation using human pulmonary fibroblasts. The cells were grown to 70 % confluency in a 12 or 24-well plate and used to test the cellular uptake of antibody-loaded PLL-coated PS by IHC and Nile Red staining. We used AF 488-labeled anti-human IgG to track the uptake by IHC and flow cytometry. MTT assay was performed to assess cytotoxicity. Untreated cells, cells treated with blank PS, cells treated with antibody alone, and cells treated with PS without PLL coating were set up as a comparison group. Cells were treated with 250ug of Ab-loaded PS (PLL-coated or uncoated) or blank PLL-PS or with 5ug of AF 488 IgG antibody overnight and were either harvested for flow cytometry or fixed with 4%PFA for IHC analysis.

### Flow cytometry analysis

Cells were trypsinized, washed, and resuspended in PBS. Healthy, untreated cells were used to establish gating parameters. Antibody uptake was quantified as the percentage of AF488? cells within each treatment group, representing successful internalization of antibody-loaded polymersomes.

### Live Cell Quantification

Cells were trypsinized, washed, and resuspended in PBS, followed by incubation with DAPI (1:3000) for 30 minutes at room temperature. Flow cytometry was performed to quantify DAPI exclusion as a measure of cell viability. Healthy, untreated cells were used to establish gating parameters. The percentage of DAPI-negative cells was reported as the percentage of live cells.

### Nile Red Staining

Cells were fixed with 4% PFA for 15 minutes at room temperature and then stained with 500nm of Nile Red for 5 minutes, followed by washing with PBS to visualize the distribution of the antibody in the cytosolic component of the cells.

### MTT Assay

MTT assay was performed to assess the acute cytotoxicity of the formulation. Briefly, pulmonary fibroblast cells were grown to full confluency in a 24-well plate. 1μL of 12-mM MTT solution was added to 100μL of media. Cells were incubated in this medium for 4 hrs. Following this step, the media was removed and 100μL of DMSO was added per well, and cells were incubated for 10 minutes at 37 °C in an incubator. After 10 minutes, the OD was taken at 540nm. After taking the readouts, cells were trypsinized and counted using bromophenol blue to account for the percentage of live cells and for normalizing the OD as a measure of viability. Healthy, untreated cells served as experimental control.

### IL-1β Assay

Culture supernatants were centrifuged at 2000 × g for 10 min at 4 °C to remove cellular debris. Cytokine concentrations in cell culture supernatants and animal serum were analyzed using the IL-1β ELISA kit (Invitrogen, DY201-05) according to the manufacturer’s protocol.

### Animals

Male,12-week-old C57BL/6 mice were used for the study. This study was approved by the Institutional Animal Care and Use Committee (IACUC) at Clemson University (Animal Use Protocol number 2025-0090). The study was carried out in compliance with the ARRIVE guidelines. Animals were acclimatized for three days before starting the study and were maintained on a standard rodent diet.

### IVIS Imaging for Acute Biodistribution

10 ml of 10mg/ml of AF488 IgG antibody loaded PLL-PS, or saline was administered via aerosol to (n=3 per group, animal use protocol number-2025-0090). 2 hours post-aerosol administration, animals were euthanized and perfused with normal saline and lungs were harvested to visualize distribution via IVIS Spectrum Fluorescence and Bioluminescence Scanner (PerkinElmer, Inc., Hopkinton, MA) using epi-illumination and an excitation and emission wavelength of 495 and 519 nm, respectively, for AF488 detection.Post imaging, lungs were formalin fixed and processed for cryo-sectioning.

### Immuno-Histochemistry –

7μm thick sections of lungs were made using a cryostat. The sections were fixed in cold acetone for 10 minutes, followed by rehydration in Dulbecco’s phosphate-buffered saline (PBS) for 15 minutes at room temperature. The sections were then washed in PBS and subsequently blocked with Background Buster (Innovex biosciences) for 30 minutes at room temperature and incubated at 4?C overnight with primary antibody for Cleaved Caspase-3. Stained sections were then incubated with appropriate secondary antibodies conjugated with either anti-rabbit Cy5 IgG for visualizing the target proteins. Finally, the aortic sections were counterstained with DAPI and mounted with an aqueous mounting medium for imaging. ImageJ software was used for image analysis.

